# The creation of functioning gastrojejunostomy following magnetic compression

**DOI:** 10.1101/2021.08.04.455102

**Authors:** Wei Qiao, Yi Lv, Zhaoqing Du, Hui Li, Lixue Du, Haitian Hu, Dichen Li

**Affiliations:** Department of Hepatobiliary Surgery, Shaanxi Provincial People’s Hospital, Xi’an, Shaanxi Province, China; National Local Joint Engineering Research Center for Precision Surgery & Regenerative Medicine, Shaanxi Provincial Center for Regenerative Medicine and Surgical Engineering, First Affiliated Hospital of Xi’an Jiaotong University. Xi’an, Shaanxi Province, China; State key laboratory for manufacturing systems engineering, Rapid manufacturing research center of Shaanxi Province, Xi’an Jiaotong University, Xi’an, Shaanxi Province, China

**Keywords:** magnetic compression anastomosis, gastrojejunostomy, anastomotic stenosis, pyloric ligation, paclitaxel-loaded

## Abstract

**Background:** Magnetic compression for creating gastrojejunostomy has many advantages according to previous studies. However, following mechanical device release after healing, the anastomotic stenosis becomes the pivotal point.

**Methods:** Rectangle-shaped magnets were used for magnetic compression in rabbits. Both paclitaxel-loaded magnets and a strategy of pyloric ligation were chosen to improve the gastrojejunostomy. Based on these choices, the half-capsule was applied to occlude the pylorus after anastomotic formation. The size and patency of the anastomoses were analyzed to evaluate the efficacy of these approaches. A histological examination was also performed.

**Results:** The positive effect of ligating the pylorus on gastrojejunostomy was significantly greater than that achieved using paclitaxel-loaded magnets during either short- or long-term follow-up. There were fewer scar tissue and collagen fibers at the anastomotic site in the treatment group than in the control group. The anastomotic aperture was of great interest at 9 months after the ligation of the pylorus following magnetic compression. In the view of the jejunum, although the aperture was barely visible, gastric juice was continuously spilling through it like a spring, and the aperture was clearly visible from the stomach side. All half-capsules failed to block the pylorus.

**Conclusion:** The effect of paclitaxel on maintaining gastrojejunostomy patency was temporary. The ligation of the pylorus ensured the long-term patency of gastrojejunostomy, and the aperture was comparable to the pylorus which could play an anti-reflux role. Further studies for the sort of gastrointestinal aperture are being planned.

## Introduction

Magnetic compression anastomosis (MCA) is a promising mechanical method for digestive reconstruction. In 1978, Kanshin and colleagues[1] reported that steady compression by magnets could be utilized to establish sutureless gastro-duodenal and ceco-jejunal anastomosis and provided apparent advantages over traditional manual sutures. Over the past 40 years, this technique has been showed to have a large number of advantages, including its convenience in use, time saving, less invasive nature, and cost-effectiveness. In addition to the speed and certainty with which an anastomosis can be made, MCA can also provide progressive compression, thus contributing to necrosis of the interposed tissues by increasing strength beyond other mechanical anastomoses[2], such as Murphy’s button and the Valtrac Ring, which have represented a breakthrough in the area of anastomotic procedures in the more than 100 years since 1892[3, 4].

The magnets maintain a permanent pressure at the anastomotic point until their release and improve the performance of circumferential airtightness so that anastomotic fistulas and bleeding can be avoided in the early stage. Following mechanical device release after healing, however, the contraction and stenosis of the stoma are the pivotal points, upon which the utility of the devices rests[5–8]. In the prior study published from our group, anastomotic contraction occurred immediately after the removal of compression magnets, regardless of whether rectangle- or cylinder-shaped magnets were used[9]. The character of contraction recovers the rhythm of dietary excretion after the reconstruction of the alimentary tract and, to some extent, can be effectively utilized to prevent digestive reflux. However, anastomotic stenosis may result in considerable morbidity or even mortality. During this process, the proliferation of scar tissue plays a critical role.

The present study investigated a solution for stenosis of the gastroenteric stoma after magnetic compression in rabbits. Base on the aforementioned research, the following issues were explored: (1) the effect of the closure of the initial outflow tract on maintaining functioning gastroenteric stoma, (2) the relationship of the primary size of the anastomosis is to its permanent patency, and (3) whether local paclitaxel-loaded magnets can suppress the production of collagen fibers in the early stage to maintain long-term patency.

## Methods

The committee on the Ethics of Animal Experiments at Xi’an Jiaotong University approved the protocol (No. XJTULAC2017-1010). The animals used in our experiment received humane treatment in strict accordance with the Animal Care and Use Committee of Xi’an Jiaotong University. All surgeries were performed under sodium pentobarbital anesthesia, and an effort was made to minimize suffering.

### Experimental animals

Japanese big ear white rabbits (n=28, 50% male) weighing 2.5 to 3.5 kg, were obtained from the Laboratory Animal Center of Xi’an Jiaotong University and housed under controlled conditions, including a 12-h light-dark cycle and a temperature of 20 ± 2°C with free access to foot and water. Before the experiment, the rabbits were adaptively fed for one week.

The animals were randomly divided into 5 groups: the initial diameter of the anastomoses was measured after MCA (n=4); MCA was performed using paclitaxel-loaded magnets (MCA-PTX, n=8); MCA was followed by the ligation of the pylorus (MCA-LP, n=8); MCA was followed by a sham operation (control group, n=5); and the magnet device for compression was connected to a half-capsule (MCA-HC, n=3).

### Design of the magnets

Rectangle-shaped magnets were used for magnetic compression to establish gastroenteric anastomosis with shells of polylactide. Their length, width and height were 16 mm, 5 mm and 2.5 mm, respectively. The magnets were magnetized perpendicular to their stress surface.

In the present study, an MCA device incorporated a pair of magnets called the gastric magnet and jejunal magnet according to their location. In the MCA-PTX group, the magnet device was wrapped in paclitaxel-loaded poly(lactic-co-glycolic acid), approximately 9 mg paclitaxel per device. In the MCA-HC group, the jejunal magnet was connected to a silk suture that was 25 cm in length. The other end of the line was tethered to a polythene tube device through a pore in a half-capsule, which was composed of polylactic acid and was 10 mm in length and 8 mm in diameter. The line was twined around the tube device and eventually encased in the half-capsule when it was introduced into the stomach. According to the design, when the MCA device fell off the anastomotic site, the half-capsule would block the pylorus due to the traction of the connecting line.

### General procedure

Anesthesia administered as intravenous pentobarbital (30 mg/kg) was followed by fixation on the operating table in the supine position with a straight neck. In the present experiment, the magnets and half-capsules were sequentially introduced into the stomach along a ferromagnetic guidewire, as described in the previous experiment[9]. Then, the jejunal magnet was squeezed into the jejunum through the pylorus. Once at the intended position, the jejunal magnet combined with the gastric magnet in the body of the stomach. This operation complied with surgical sterile standards. After the rabbits recovered from anesthesia, they were returned to their cages and allowed free access to food and water.

An abdominal X-ray was taken immediately to mark the initial position of the magnet device as the base level, and this data was used to confirm whether the device became unexpectedly separated or fell off the anastomotic stoma during the experimental period.

### Specimen collection

When the endpoint was achieved in the experiment, the animals were euthanized with an excess of pentobarbital. The anastomosis was immediately harvested, and its diameter was measured in the natural state. Finally, both hematoxylin-eosin and Masson’s trichrome staining were performed to observe histomorphology by optical microscopy.

### Statistical analysis

The diameter of the anastomotic stoma was measured in millimeters (mm) and was presented as the average of the long and short diameter. The force per square centimeter exerted at the magnet device surface was shown as the associated pressure. The quantitative data were expressed as the mean ± standard deviation. Student’s *t*-test and one-way ANOVA were used in SPSS 22 for statistical analysis. *P* < 0.05 was considered to indicate statistical significance. The sex of the animals was not considered a factor in this study.

## Results

In this study, all animals survived to the endpoint. There was no intestinal obstruction, fistula, or infection in any of the groups. Each incision healed well. When the magnet pairs fell off, they were freely extruded from the anus. The formation time of the anastomotic stoma was 12.1 ± 1.4 days following the MCA procedure (n=20) and 11.4 ± 0.9 days after the procedure when paclitaxel-loaded magnets were used (n=8, *P* > 0.05).

In the test of MCA-HC, all three half-capsules failed to act as expected to block the pylorus, and they were extruded from the pylorus before the magnet devices released (**Fig 1**). These rabbits were randomly transferred into the group of measuring initial diameter, the control group, or the MCA-LP group, respectively.

**Fig 1.**
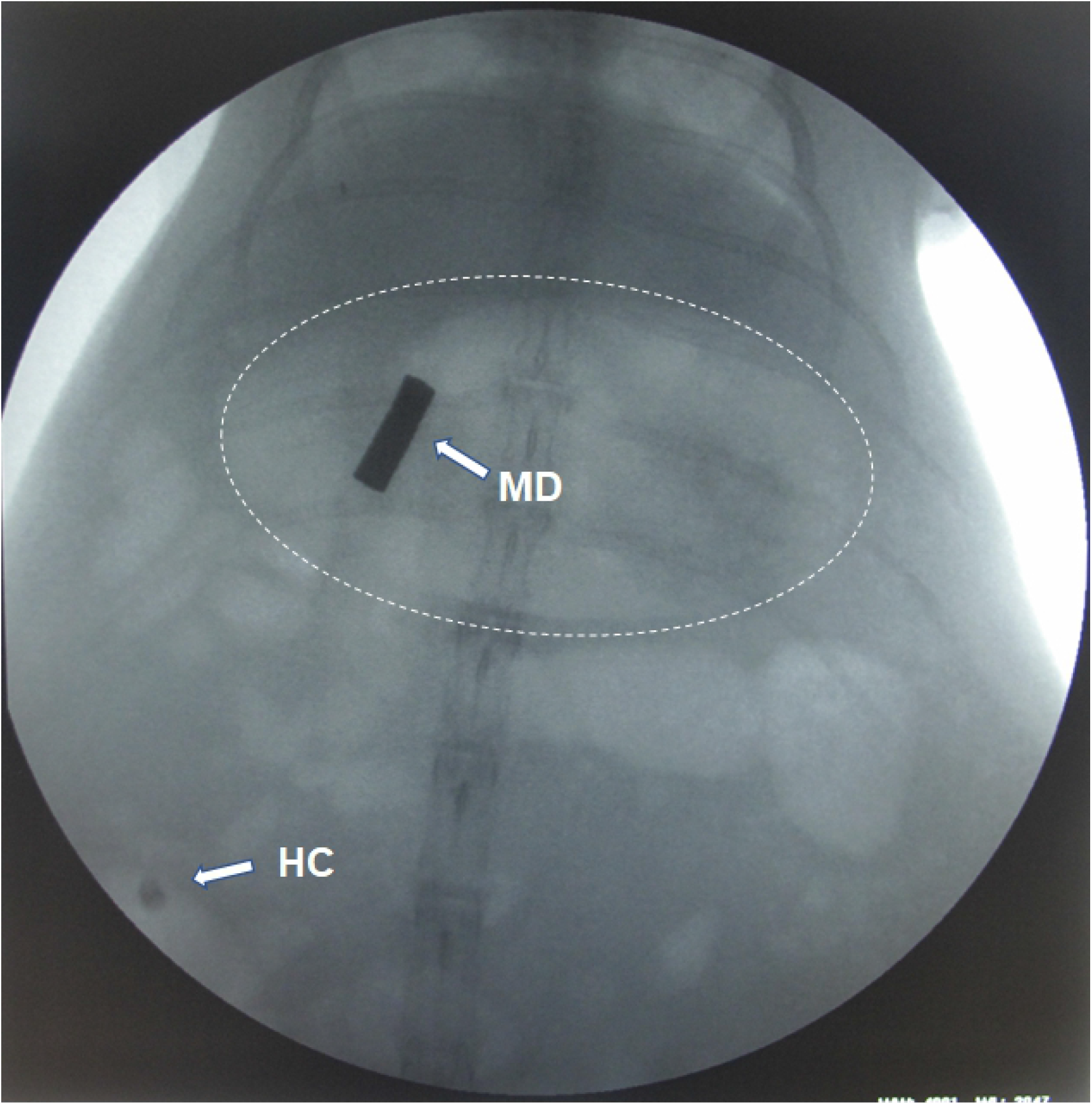
Abdominal X-ray examination at 2 weeks after magnetic compression in the MCA-HC group. The half-capsule (**HC**) has been extruded from pylorus into the bowel and the magnet device (**MD**) was still at the site of gastrojejunostomy. The dotted oval indicates the contour of the stomach.

The abdominal adhesion was barely visible, and the diameter of the jejunal loop with the gastrojejunostomy significantly increased in each group. The initial gastrointestinal anastomosis was 5.0 ± 0.5 mm in diameter (**Fig 2A**), and the calculated area was 19.6 mm^2^, which was not in accordance with the expected value of 80 mm^2^ given a relative contraction of 75.5%.

**Fig 2.**
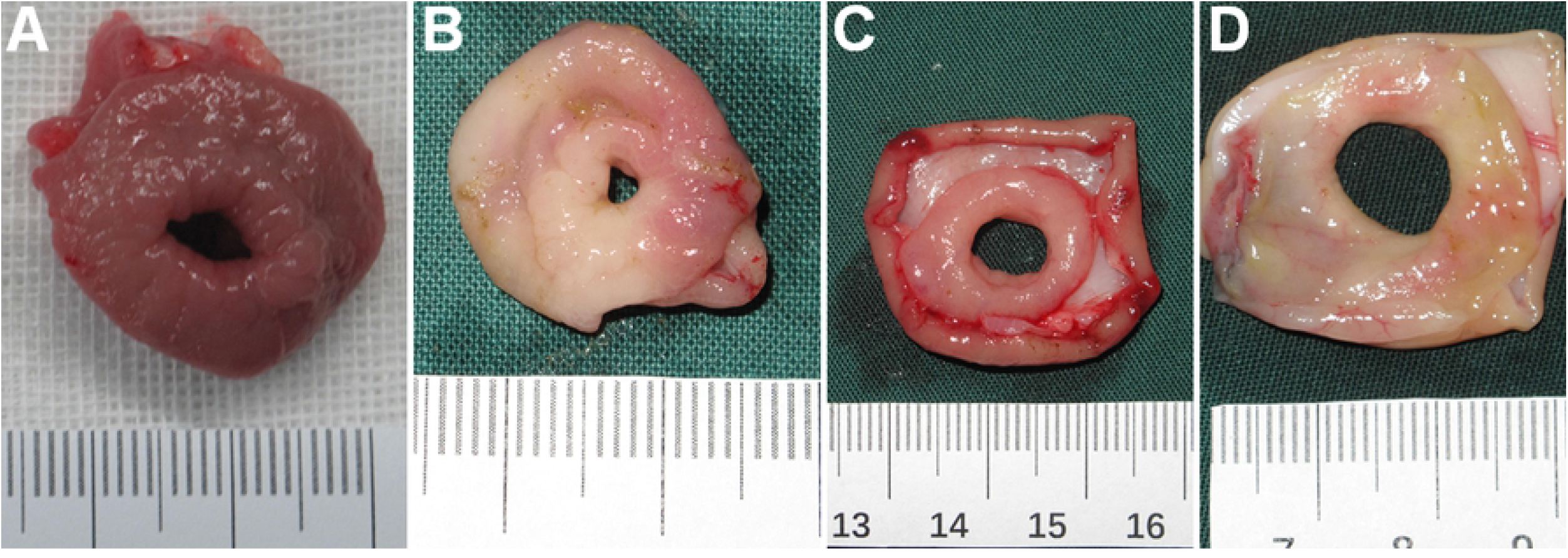
Gross appearance of gastrojejunostomy. (A) The initial anastomosis. The anastomoses at 50 days after establishment (B, in control) using magnets followed by the sham operation, (C) using paclitaxel-loaded magnets and (D) using magnets followed by the ligation of the pylorus.

The anastomotic diameter was reduced to 2.2 ± 0.2 mm at 50 days after the MCA procedure (*P* < 0.01, *vs*. the initial diameter; **Fig 2B**). In the present study, a diameter of 7.1 ± 0.2 mm was measured at 50 days after the operation in the MCA-PTX group (*P* < 0.01 *vs*. control group; **Fig 2C**), while the diameter was 8.5 ± 0.4 mm in the MCA-LP group (*P* < 0.01 *vs*. control group; *P* < 0.01 *vs*. MCA-PTX group; **Fig 2D**).

The diameter of the anastomotic stoma was 5.0 ± 0.3 mm at 9 months after MCA-LP (*P* > 0.05 *vs*. the initial diameter; *P* < 0.01 *vs*. the diameter at 50 days after MCA-LP; **Fig 3**). The aperture of anastomosis was of great interest at 9 months after the procedure in the MCA-LP group. On the outside view, the side-to-side anastomoses between the stomach and jejunum were complete (**Fig 3A**). In a window sectioning the jejunum, although the aperture was barely visible, gastric juice was continuously spilling through it like a mountain spring (**Fig 3B**), and the aperture was clearly visible from the stomach side (**Fig 3C**). The gastrojejunostomy was covered by mucosa with fibrosis in the submucosal and muscular layers. There was less scar tissue and fewer collagen fibers at the anastomotic site in the experimental group than in the control group (**Fig 4**).

**Fig 3.**
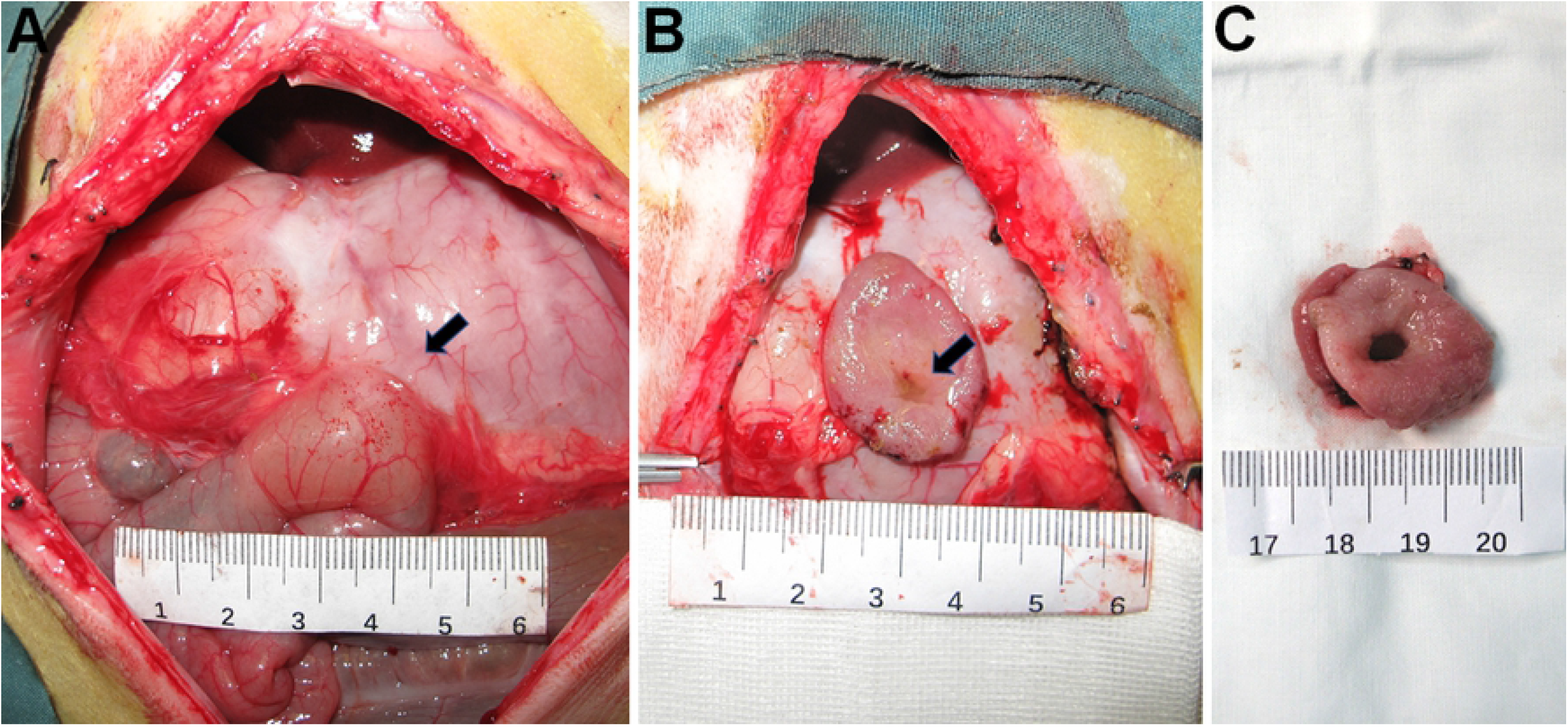
The aperture of the anastomosis at 9 months after the procedure in the MCA-LP group. (A) Outside view showing the side-to-side anastomosis between the stomach and jejunum. (B) Section through the jejunum showing that the aperture was barely visible, but the gastric juice was continuously spilling through it like a mountain spring. (C) The aperture from the stomach side. The black arrow points to the anastomotic site.

**Fig 4.**
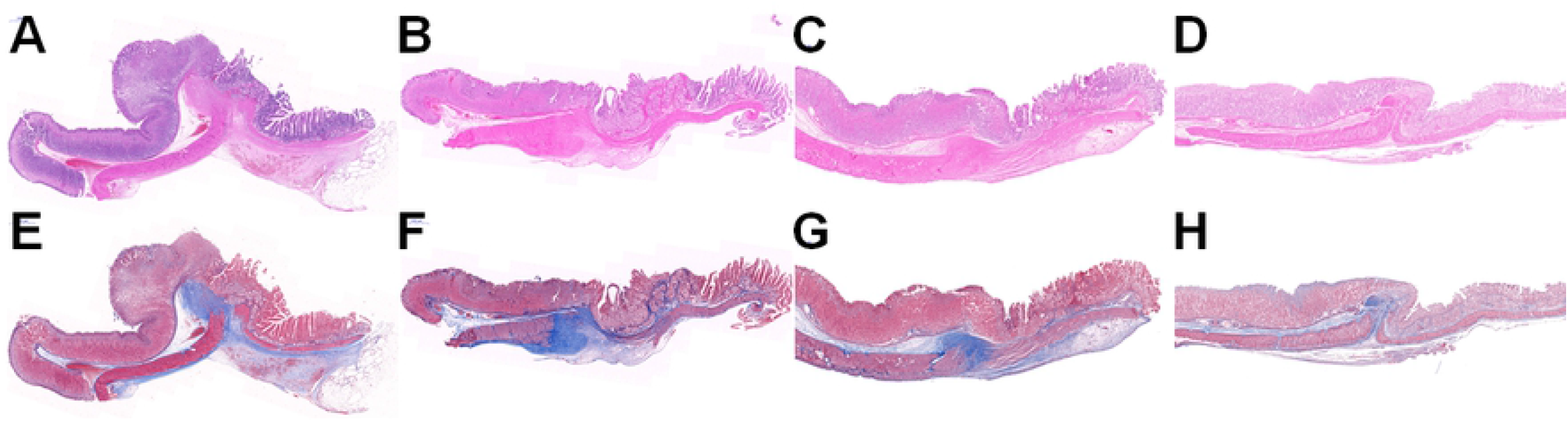
Histological sections of gastrojejunostomy. (A, E) The initial anastomosis. The anastomoses at 50 days after establishment (B, F) using magnets followed by the sham operation, (C, G) using paclitaxel-loaded magnets and (D, H) using magnets followed by the ligation of the pylorus. (Top: hematoxylin-eosin staining. Bottom: Masson staining)

Nevertheless, the bypass established between the stomach and jejunum was closed at 9 months after the procedure in the MCA-PTX group and was not covered by mucosa. The results of both gross examination and histological sections were shown in **Fig 5**.

**Fig 5.**
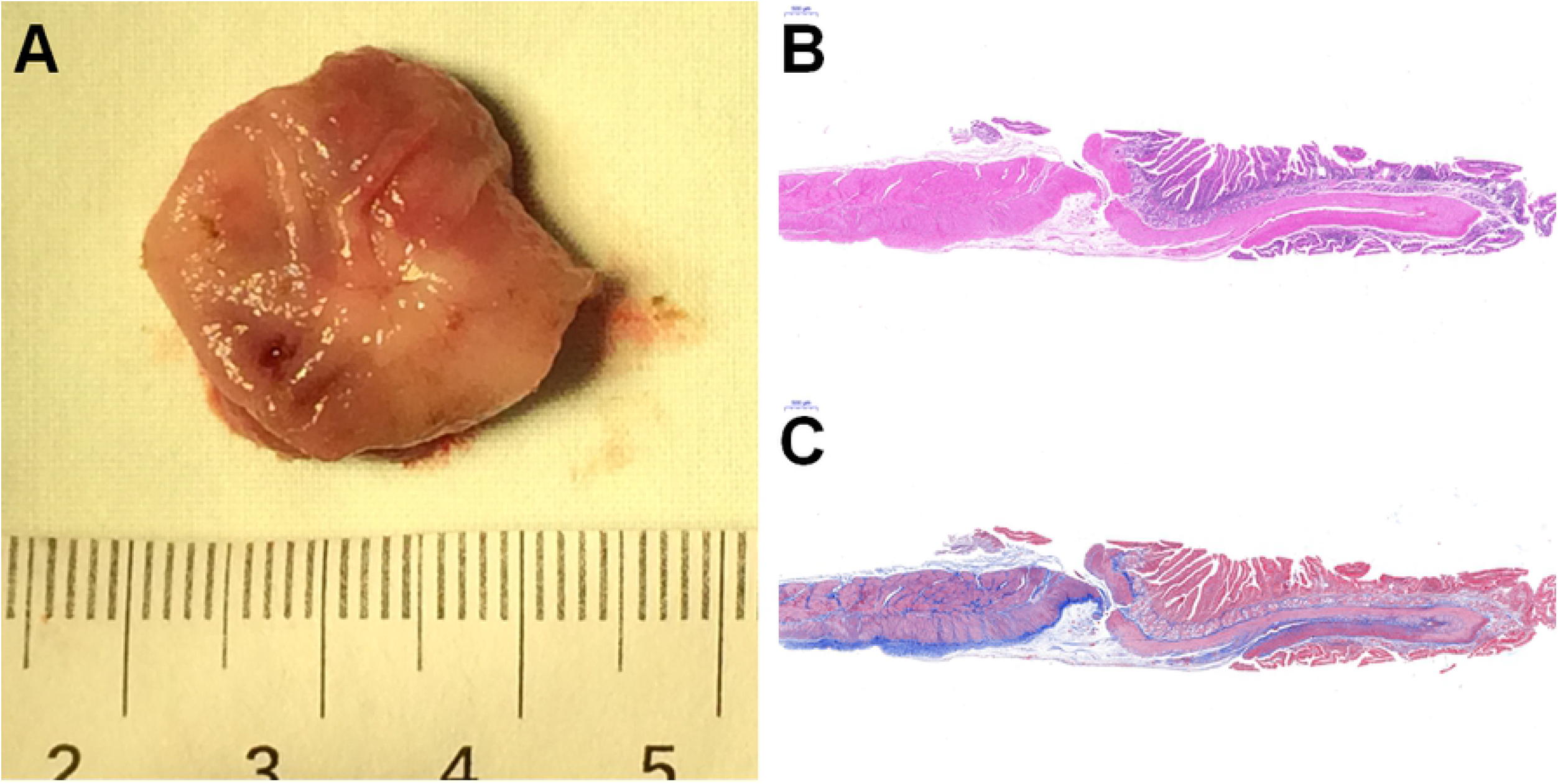
The closed gastrojejunostomy at 9 months after the procedure in the MCA-PTX group. (A) Gross appearance. Histological examination stained with hematoxylin-eosin (B) and Masson staining (C).

The magnets firmly attracted to each other with no accidental separation. The compression force applied by the paired magnets was 9.1 ± 0.4 N, and the associated pressure was 11.4 ± 0.5 N/cm^2^ with no spacing. Considering the thickness of the interposed tissue, the force was 4.4 ± 0.9 N with an associated pressure of 5.6 ± 1.0 N/cm^2^ according to the distance of 1 mm.

## Discussion

It is generally believed that both fibroblast proliferation and the synthesis of collagen fiber play important roles in the process of anastomosis establishment. Nevertheless, excessive collagen accumulation causes pathological scar formation during wound healing and acts as a critical factor leading to an anastomotic stricture. Paclitaxel has been shown to have the anti-fibrotic effect[10]. In the MCA-PTX group, although the effect of paclitaxel on the synthesis of collagen fiber was negative, it neither postponed the formation of gastrojejunostomy nor resulted in the anastomotic leakage. These results indicated that the experimental group showed apparent advantages on gross and histologic examination. Compared with the control group, the experimental group had less scar tissue, which has been shown to be helpful in reducing the incidence of anastomotic stenosis[11, 12].

Both fibrosis and inflammation are critical to healing and are likely beneficial until creating a stenosis. In traditional approaches, the scar gradually forms during the inflammation phase, approximately one month after the operation[13]. In this study, the first observation date was set at the 50 days after the operation. In the control group, the aperture had contracted to less than one-half of its original diameter. In the previous study that used smaller magnets (7.2×3.5×2.0 mm), the initial diameter of gastrojejunostomy was 1.5 mm, and the gastrojejunostomy was completely closed within 12 days (Qiao W, unpublished data). The primary size of the anastomosis is related to its permanent patency.

The local administration of drugs such as paclitaxel-loaded magnets, can improve the patency of the anastomotic stoma in the early stage, but its long-term effect was not substantial. The positive effect of ligating the original outflow tract-pylorus on re-established gastrojejunostomy was significantly greater than the local effect of paclitaxel during either short- (at 50 days) or long-term (at 9 months) follow-up. After the establishment of gastrojejunostomy by magnetic compression, the timely closure of the original tract can avoid anastomotic stricture.

The type of gastrointestinal aperture observed at 9 months after the procedure in the MCA-LP group was comparable to the pylorus which could play an anti-reflux role. This finding is interesting enough that surgeons should re-evaluate the advantages of magnetic compression anastomosis. The limitation of this part of the study was the relatively small sample size.

In addition, the pressure produced by the magnets used in this study was the minimum reported in all studies of successful gastrojejunostomy[13, 14]. This investigation may facilitate a factor of robustness in the design of future devices. It should be noted that the pressure between mating magnets varies with the thickness of the interposed tissue, which evolves over time during the process of magnetic compression.

In the MCA-HC group, all three half-capsules passed through the pylorus before the magnet devices fell off the anastomoses, and no half-capsule was found in the stomach. These findings indicate that the half-capsules used in this trial were not suitable for the occlusion of the pylorus, but the novel strategy for occluding the pylorus after anastomotic formation is still worth further investigation.

## Conclusions

The magnets with larger sizes are recommended to establish a stoma in the digestive tract. The effect of paclitaxel is temporary to maintain the gastrointestinal patency. The ligation of the pylorus ensures the long-term patency of gastrojejunostomy, and its aperture is comparable to the pylorus which could play an anti-reflux role. Based on this study, further studies for the sort of gastrointestinal aperture are being planned.

## Supporting information

**S1 File. Completed “The ARRIVE Guidelines Checklist”.**

